# A network-based model for drug repurposing in Rheumatoid Arthritis

**DOI:** 10.1101/335679

**Authors:** Ki-Jo Kim, Navneet Rai, Minseung Kim, Ilias Tagkopoulos

## Abstract

**Background:** The identification of drug repositioning targets through computational methods has the potential to provide a fast, inexpensive alternative to traditional drug discovery process. Diseases where complicated pathophysiology evolves over time are excellent targets for such methods. One such disease is Rheumatoid Arthritis (RA), a chronic inflammatory autoimmune disease, where the drug survival at 5-years is less than 50% for patients treated with disease-modifying anti-rheumatic drugs (DMARDs).

**Methods and Findings:** We have developed a network-based approach for drug repositioning that takes into account the human interactome network, proximity measures between drug targets and disease-associated genes, potential side-effects, genome-wide gene expression and disease modules that emerge through pertinent analysis. We found that all DMARDs, except for hydroxychloroquine (HCQ), were found to be significantly proximal to RA-related genes. Application of the method on anti-diabetic agents, statins and H2 receptor blockers identified anti-diabetic agents – gliclazide, sitagliptin and metformin – that have similar network signatures with the DMARDs. Subsequent *in-vitro* experiments on mouse fibroblast NIH-3T3 cells validated the findings and the down regulation of six key RA-related inflammatory genes. Our analysis further argues that the combination of HCQ and/or sulfasalazine with methotrexate (MTX) is predicted to have an additive synergistic effect in treatment based on network complementarity. Similarly, leflunomide and tofacitinib were found to be suitable alternatives upon chemoresistance to MTX-based double/triple therapy, given the complementary network signatures and overlapping critical target hubs.

**Conclusions:** Our results corroborate that computational methods that are based on network proximity, among other contextual information can help narrow down the drug candidates for drug repositioning, as well as support decisions for combinatorial drug treatment that is tailored to patient’s needs.

**Author summary:** The network-based proximity between drug targets and disease genes can provide novel insights on the repercussion, interplay, and reposition of drugs in the context of disease. Disease-modifying anti-rheumatic drugs (DMARDs) located significantly close to rheumatoid arthritis (RA)-associated genes and RA-relevant pathways. Three anti-diabetic agents were identified to have an anti-inflammatory effect like DMARDs. We built RA disease module encompassing the emerging small-molecule targets and functional neighbors, which better explained the RA pathophysiology. By proximity and network robustness, tofacitinib and tocilizumab were the most potent for RA disease module as a single agent, and this is consistent with clinical observation. Side effects of clinical importance are predictable by measuring network-based proximity between drug targets and side effect protein. Network-based drug-disease proximity offers a novel and clinically actionable information about drugs and opens the new possibility to drug combination and reposition.

## Introduction

Cellular state can be viewed as a well-orchestrated choreography with a touch of chaotic behavior. Genes, proteins and other chemical compounds rarely act in isolation, instead they participate in pathways and mechanisms that give rise to the various biological functions of the cellular network [1]. In the context of disease, cellular state is perturbed by the activation of common pathways through a variety of dysregulated genes and aggregated activity of compounds, leading to a similar phenotype [2, 3]. In a similar fashion, drugs exert their therapeutic effects by modulating molecular pathways and molecule(s) [4], with possible off-target effects due to the presence of system-wide mechanisms and processes [5]. Direct study of the respective pathways and biological processes is cumbersome; however, systemic analysis of large integrated datasets has recently provided opportunities to unravel novel connections, and hence bridge knowledge gaps to better understand human diseases [6, 7]. Using the proximity of the compounds and their targets to key disease genes has emerged as a possible method to identify potential compound-disease pairs. These analyses are based on the hypothesis that the closer a drug’s target is to the disease-associated network, the higher the chance it will influence the disease’s progression and state [8]. This principle, which is the foundation of this study, can guide drug exploration towards drug repurposing and personalized medicine.

Rheumatoid arthritis (RA) is a chronic inflammatory autoimmune disease that causes progressive joint destruction [9]. Dynamic, complex interactions between a multitude of environmental and genetic factors affect the disease development and activate pathogenic mechanisms, some of which are poorly understood [9]. Despite the introduction of new drugs targeting key mediators of RA – cytokines (TNF and IL-6) and immune cell regulators (B7 molecules and CD20), over 50% of patients only partially respond to therapy [10]. Moreover, disease flare and efficacy reduction is common in patients with satisfactory initial response to medication [11]. Hence, there is a current and constant need for the development of new RA drugs, which can be tailored to specific targets and personalized based on the patient profile [12].

In this study, we analyzed the relationship between disease-modifying, anti-rheumatic drugs (DMARDs) and RA-associated subnetworks using proximity measures in an unsupervised and unbiased network-based framework. First, we explored the distance of key molecules that are targeted by DMARDs to RA-associated genes. Then we focused on the pertinent pathways and respective biological processes associated based on the distance metrics with a focus on the degree the drug-perturbed pathways cover pathogenic processes of RA. Finally, we created predictors of drug effectiveness and retargeting potential based on network-based signatures, which resulted in 3 potential RA drug candidates that were further validated through in vitro screening. **Fig 1** depicts the overarching methodology for our data collection and proximity analysis performed.

**Fig 1.**
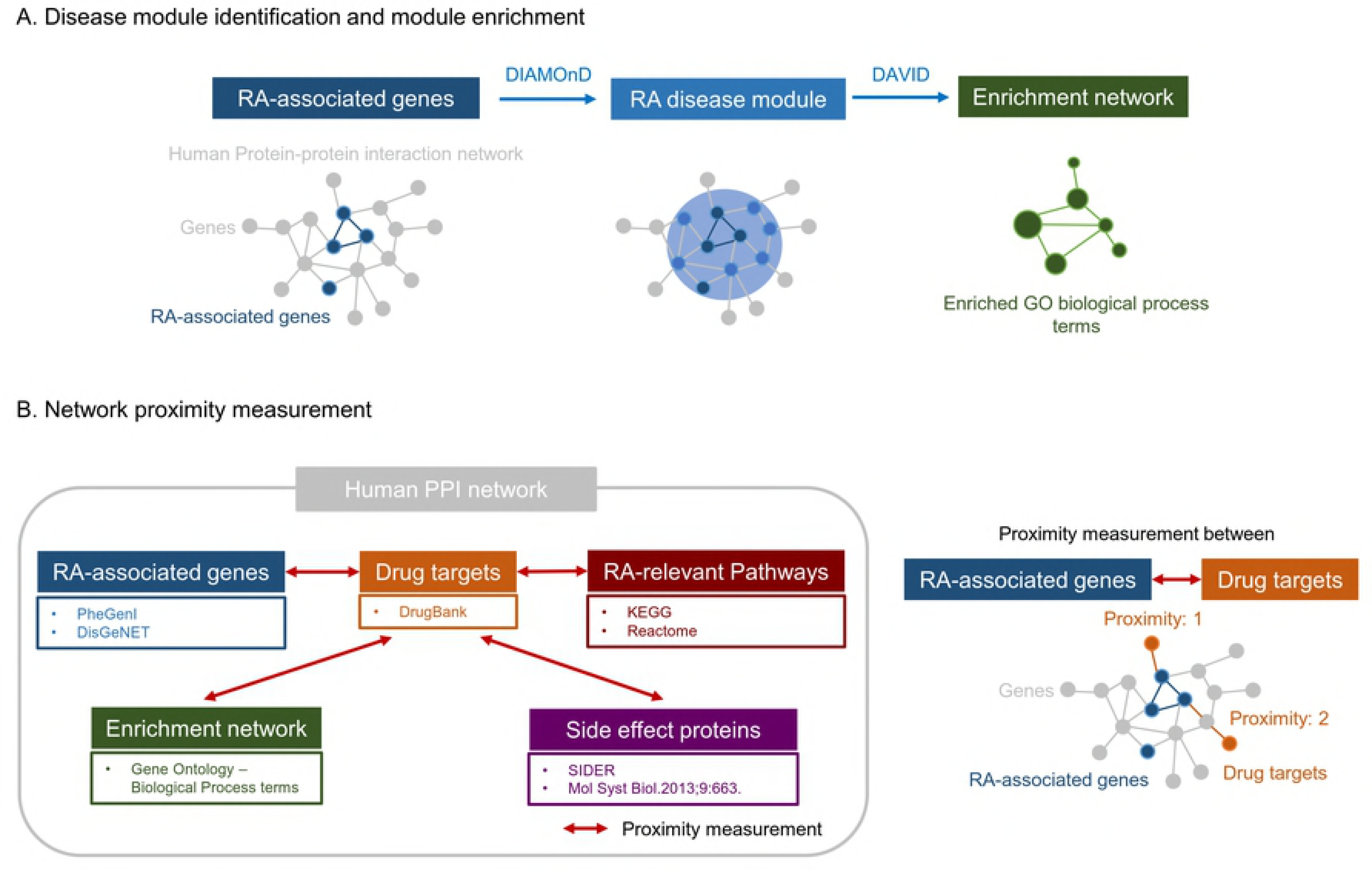
Overview of the computational approach. (**A**) The RA-associated genes are used as seed for identification of the RA disease module, by using the DIAMOnD module finding algorithm on the human interactome network. Once the RA disease module has been constructed, GO analysis is performed through DAVID for assessment of the related biological processes. (**B**) Overview of the network proximity data sources and methodology.

## Materials and Methods

### The human interactome

A previous study [13] constructed a model of the human interactome based on its protein-protein interaction (PPI) network built based on the experimentally documented human physical interactions from TRANSFAC [14], IntAct [15], MINT [16], BioGRID [17], HPRD [18], KEGG [19], BIGG [20], CORUM [21], PhosphoSitePlus [22], as well as an individually published large-scale signaling network [23]. This human interactome dataset contains 141,150 interactions among 13,329 proteins and was found to perform better in capturing the therapeutic effect of drugs as compared to functional networks from STRING [24] and other high-throughput binary screens [8]. We are using this human interactome network as our basis to conduct our proximity analysis.

### RA-associated genes and drug targets

RA-associated genes were retrieved from PheGenI [25] and DisGeNET [26]. 147 and 219 genes were found to be associated with RA in the PheGenI and the DisGeNET database, respectively. 252 unique genes belong to human interactome elements (**S1 Table**). Information on drug-target molecules were extracted from the DrugBank database (**S2 Table**) [27].

### Network-based proximity measure

To quantify the network-based relationship between drug targets and disease protein, we used the metric of *closest proximity d*_c_ that is defined as the average shortest path length between the drug’s targets and the nearest disease protein [8]. We found that the *closest proximity* metric offered superior performance when compared to other network-based distance measures such as the shortest measure, the kernel measure, the center measure, and the separation measure [8]. Briefly, given *S*, the set of disease proteins, *T*, the set of drug targets and *d(s, t)*, the shortest path length between nodes *s* and *t* in the network, the closest proximity *d*_c_ between S and T is given by:

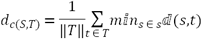

To assess the statistical significance of the closest proximity, we introduced a measure of relative proximity *z*_c_ that captures the distribution of distances between random and RA-associated proteins by calculating its z-score. The reference distance distribution was generated by calculating the distance between RA-associated proteins and the randomly selected group of proteins matching the size and the degrees of the drug targets in the network, re-calculated over 1,000 random runs. As such, the relative proximity *z*_c_ is given by:

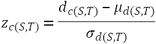

where *μ*_*d*(*S,T*)_ and *σ*_*d*(*S,T*)_ are the mean and standard deviation of the reference distribution. If *z*_c_ ≤ – 0.15, a drug was considered as proximal to the disease. To select the threshold we performed parameter sweeps (grid-search) and also investigated the sensitivity of the final results based on the threshold neighborhood. We observed that for thresholds close to –0.15, both sensitivity and specificity have optimal values, drug-disease association coverage is in agreement with expert opinion and known knowledge and that interactome-based proximity is not changed significantly by repeating the analyses by using either National Drug Files or the KEGG database, in agreement with a previous study [8].

### Drugs used in this study

The DMARD drug class here includes the following drugs: methotrexate (MTX), hydroxychloroquine (HCQ), sulfasalazine (SSZ), leflunomide (LFNM), tofacitinib (TFCN), adalimumab (ADLM), etanercept (ETAN), infliximab (IFXM), tocilizumab (TCZM), rituximab (RTXM), and abatacept (ABCT). For comparison, selected as controls were some of the anti-diabetic agents (ADA), statins, H2 receptor blockers (H2RB): metformin (MFOM), gliclazide (GCZ), glimepiride (GMPD), pioglitazone (PGTZ), sitagliptin (STGT), canagliflozin (CNGZ), exenatide (EXNT), simvastatin (SMVT), atorvastatin (ATVT), pravastatin (PRVT), fluvastatin (FLVT), rosuvastatin (RSVT), cimetidine (CMTD), and ranitidine (RNTD).

### Network-based pathway and side-effect proximity analysis

To identify the biological pathways affected by a drug in the human interactome, we used the closest distance measure to assess the proximity between drugs and pathways. The drug-pathway proximity is the normalized distance calculated between the drug targets and proteins belonging to a given pathway. Similar to our calculation of drug-disease proximity, 1000 randomly selected protein sets matching the original protein sets in size and network degree were used to calculate the mean and the standard deviation of the *z*-score. We used gene sets of all KEGG [28] and Reactome pathways [29] provided in MSigDB [30].

To check if a drug was adjacent to the proteins provoking certain side effects, we calculated the network-based proximity of drug targets to side-effect proteins (same for disease and pathway proteins). Side-effect proteins were referred to the result by a Fisher’s exact test-based enrichment analysis [31] and 99 proteins and 230 side effects were paired. Drug-specific side effects were imported from the SIDER database [32]. Since there is redundancy in the types of side effects for different proteins, if multiple proteins were candidates for drug-associated side effects, a protein more proximal to the drug target was selected.

### Disease Module Detection Method

To obtain the RA-specific disease module in the human interactome, we used the Disease Module Detection (DIAMOnD) algorithm [33, 34], which is robust over a wide range of noise and superior performance as compared to the other methods. This algorithm is based on the observation that potential disease genes have a propensity to interact with the known disease genes and prioritizes the proteins in the network based on their topological proximity to seed proteins [13]. Briefly, if there is a number of *s* seed proteins associated to a certain disease in the network, the probability for a protein with a total of *k* links to have exactly *k*_*s*_ links to seed proteins is given by the hypergeometric distribution:

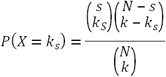

The connectivity *p-*value, i.e. the cumulative probability for the observed or any higher number of connections is given by:

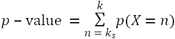

The protein with the highest rank (i.e. lowest *P* value) is added to the basic set of seed genes and the gene encoding the protein added by the algorithm are designated as ‘DIAMOnD gene’. In the next iteration, the updated set of seed genes is used to identify the highest ranking protein, until the entire network is completed. The analysis identified statistically significant enrichment for RA-relevant genes among the identified DIAMOnD genes. Since there was no significant further gain beyond 550 iteration steps, we considered the first 550 DIAMOnD genes to have the most significant RA association (**S3 Table**). The module that starts at 85 directly connected seed genes was added by 550 DIAMOnD genes and was further connected to a number of isolated seed genes (103 genes) through DIAMOnD genes. As a result, a putative RA disease module of 738 genes in total was created.

### Validation of DIAMOnD genes

To determine the cutoff size of DIAMOnD genes, we used four RA-relevant validation data – RA gene expression data, gene sets from KEGG, Reactome pathways, and GO biological process terms. For RA gene expression data, we searched the NCBI Gene Expression Omnibus (GEO, January 2017, http://www.ncbi.nlm.nih.gov/geo/) and ArrayExpress database (https://www.ebi.ac.uk/arrayexpress/) for all gene expression dataset using ‘rheumatoid arthritis’ and ‘synovial tissue’ as a search term, resulting in 14 public available human microarray data. Since there was a trade-off between the number of studies to include and the number of genes that are within the intersection from all datasets, we optimized the product of the two by selecting the point where these two trends cross. We finally selected 11 expression data with GEO series (GSE) ids: GSE77298, GSE48780, GSE45867, GSE36700, GSE24742, GSE15602, GSE21537, GSE12021, GSE55584, GSE55457, GSE55235 and the united collection included the 256 RA, 41 osteoarthritis, and 36 normal samples. After normalizing and removing the batch effect, we performed differentially expression analysis and selected the gene candidates that were identified in three independent methods (empirical Bayes, significance analysis of microarray, and ranked product) and finally filtered the 2212 shared up-regulated genes. The gene sets of KEGG, Reactome pathways or GO terms that are significantly enriched within the given set of seed genes were obtained using the Database for Annotation, Visualization, and Integrated Discovery (DAVID) version 6.8 software and Reactome Analysis tool [29, 35] (*P* value < 0.05). For each DIAMOnD gene, it is considered as a “hit” if it has at least one annotation that is in the gene sets significantly enriched within the seed genes (Fisher’s exact test, *P* value < 0.05). To compensate for the dependence of *P* values on the underlying set size, we used a sliding-window approach: At each iteration step i, we consider all DIAMOnD genes in the interval [i – 252/2, i + 252/2], thereby obtaining sets of the same size as the seed genes that can be compared to each other.

### Assessment of network robustness

To assess the robustness of the networks against damage, we used the iterative ‘random’ and ‘targeted’ removal of nodes [36, 37]. Nodes were deleted randomly from the graph irrespective of the degree of the node. Or only the nodes targeted by the drugs were selectively removed. After each deletion, we computed the evolution of the size of the largest connected component (LCC), which means minimum path length under a random or targeted attack of a graph. This can tell us how much the drug can disrupt the network of disease module.

### Functional enrichment analysis

We performed functional enrichment analysis focusing on the genes of interest using DAVID software [35]. Terms were regarded significant if the *P* value (EASE score) is lower than 0.01, the gene count more than 3, and the fold enrichment larger than 1.5. The enrichment results were visualized with the Enrichment Map format, where nodes represent gene-sets and weighted links between the nodes represent an overlap score depending on the number of genes two gene-sets share (Jaccard coefficient) [38]. To intuitively identify redundancies between gene sets, the nodes were connected if their contents overlap by more than 10%.

### Single sample gene-set enrichment analysis

To test for gene enrichment in individual samples from RA patients (rather than in a group of samples from RA subjects as the former is clinically more relevant), we used a single sample version of gene-set enrichment analysis (ssGSEA), which defines an enrichment score as the degree of absolute enrichment of a gene set in each sample within a given data set [39]. The gene expression values for a given sample were rank-normalized, and an enrichment score was produced using the Empirical Cumulative Distribution Functions (ECDF) of the genes in the signature and the remaining genes. This procedure is similar to the GSEA technique, but the list is ranked by absolute expression in one sample.

### In-vitro validation

#### Materials

The mouse fibroblast NIH-3T3 cell line was used throughout the experiments with the Dulbecco’s Modified Eagle’s Medium with L-Glutamine, 4.5g/L Glucose and sodium pyruvate (Corning Inc.), HyClone Newborn Bovine Calf Serum (GE Healthcare), Gliclazide and Sulfasalazine (ACROS Organics), Metformin hydrochloride (Mp Biomedicals Inc.), Cimetidine (Alfa Aesar), Sitagliptin phosphate (Biotang Inc.), Penicillin/Streptomycin and Methotrexate hydrate (Sigma). The RevertAid First strand cDNA Synthesis kit, VeriQuest SYBR Green qPCR Master Mix, and Nunc EasYFlask 75cm^2^ were purchased from Thermo Scientific.

#### Cell culture and treatment

The expression of RA related inflammatory genes (IL-6, IL-1β, MCP-1, RANTES, CXCL1, and IκB) was measured on the NIH-3T3. The cells were treated with following immunomodulatory agents: gliclazide (10 µg/mL), metformin (2 mM), sitagliptin (0.1 mM), methotrexate (10 µM), sulfasalazine (1 µM). cimetidine (10 µg/mL) and TNF-α (10 ng/mL). Gene expression in untreated NIH-3T3 cells was used as the baseline. Two different strategies were used to measure the efficacy of these drugs on the inflammatory genes. In first strategy, NIH-3T3 cells were treated with above mentioned immunomodulatory agents for 24 h at 37^°^C in an incubator saturated with 5% CO_2_, and samples were preserved in RLT buffer (Qiagen) at −80^°^C, as described in the RNAeasy kit (Qiagen) manual. In second strategy, NIH-3T3 cells were first treated with above mentioned immunomodulatory agents, excluding TNF-α, for 2 h and then were treated with the TNF-α for 24 h at 37^°^C in an incubator saturated with 5% CO_2._ Samples were preserved for RNA-isolation as described before.

#### RNA isolation and Real Time PCR

Total RNA was isolated using RNAeasy kit (Qiagen) as described in the manual. Quantification of total RNA was done using Qubit (Life Technologies). Subsequently, random hexamer assisted first strand of cDNA was synthetized from 5 µg of total RNA using RevertAid cDNA kit. Real time gene expression was measured using Veriquest SYBR mix on ABI Viia 7 RT-PCR and primer sets described in the table below. GAPDH was used as endogenous control (reference gene). Ct values were calculated in Quant Studio Real Time PCR Software with default settings.

### Statistical analysis

For continuous distributed data, between-group comparisons were performed using the paired or unpaired *t*-test. Categorical or dichotomous variables were compared using the chi-squared test or Fisher’s exact test. All analyses were conducted in *R* (The R Project for Statistical Computing, www.r-project.org).

## Results

### Proximity between DMARDs and RA-associated genes in the interactome

The drug-target information was gathered from DrugBank database [27] (**S2 Table**) and bipartite network between RA-associated genes and DMARDs is depicted in **Fig 2A**. The relative proximity between DMARDs and RA-associated genes is plotted in **Fig 2B**. All DMARDs except for HCQ were found to be significantly proximal to RA by *z*_c_ < –0.15. ABCT was most proximal to a group of RA-associated genes (*z*_c_ = –3.09) and LFNM was least proximal (*z*_c_ = –0.22). MTX had a proximity of *z*_c_ = –1.61 though it was defined to target a single molecule, dihydrofolate reductase. In contrast, molecules targeted by statins and H_2_-receptor blockers placed much far from RA-associated genes (*z*_c_ > 0.0).

**Fig 2.**
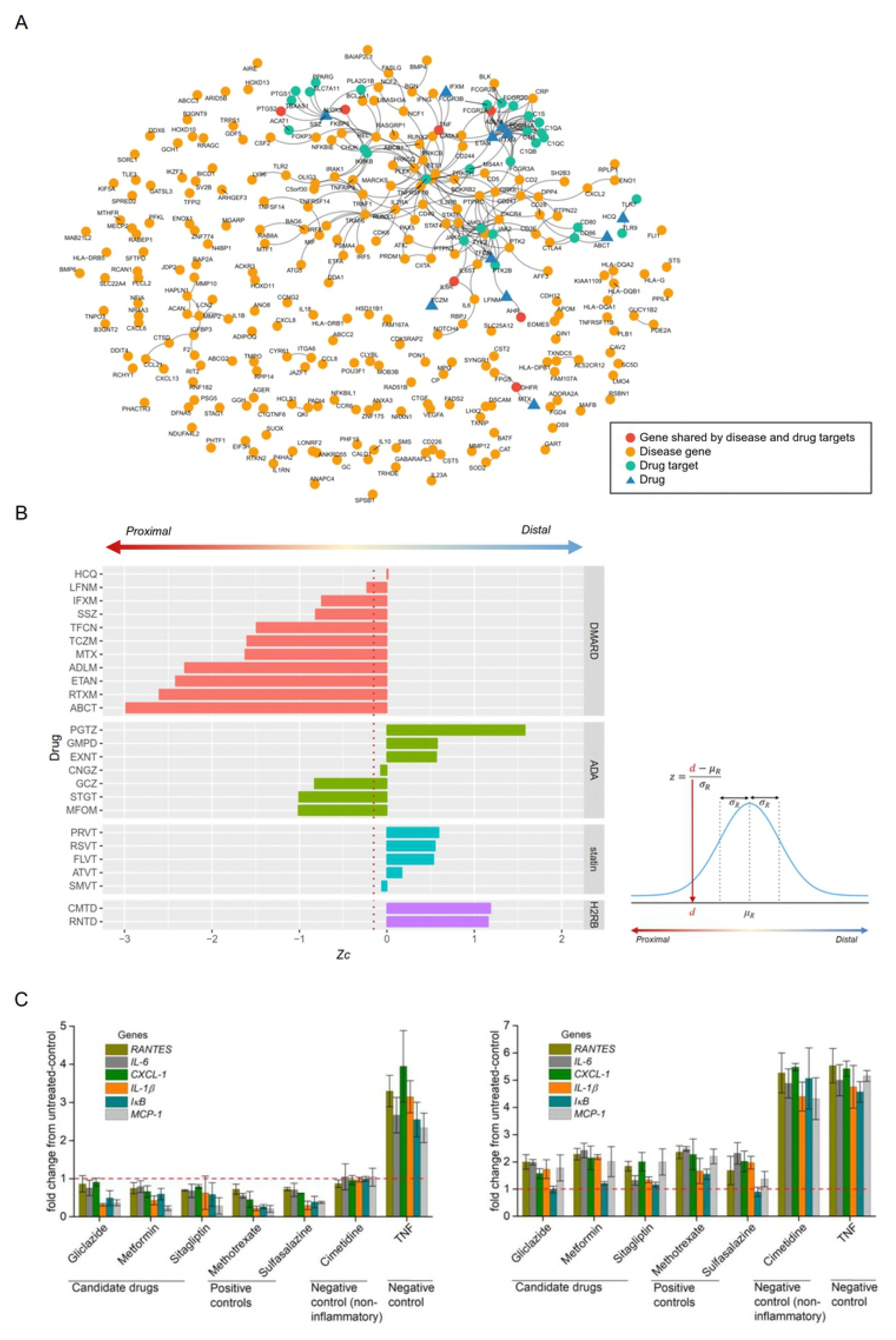
Gene-Drug target network and proximity between RA-associated genes and DMARDs. (**A**) A graphical representation of all the interactions between RA-associated genes, DMARDs, and their targets. Red nodes are RA-associated genes that are also drug targets. Yellow nodes are disease genes, green nodes are drug targets, and blue triangles are DMARDs. (**B**) Given a set of disease proteins *S*, and the set of drug targets *T*, the closest distance *d* is the shortest path length between all members of *S* and *T* in the network. The relative proximity (*z*) was calculated by comparing the distance *d* between *T* and *S* to a reference distribution of distances between RA-associated proteins and the 10^3^ groups of randomly selected proteins matching the size and the degrees of the drug targets in the network. The red dotted line corresponds to the significance threshold (*z* ≤ –0.15). All DMARDs excluding HCQ were found to be significantly proximal to RA-associated genes. From non-DMARDs, PGTZ, GMPD, EXNT, statins, and H2RB were found to not be associated to RA-related genes, while GCZ, MFOM, and STGT have DMARDs-like signatures. (**C**) GCZ, MFOM, and STGT mediated repression of RA associated inflammatory genes in mouse fibroblast NIH-3T3 cells. Responses of inflammatory genes after 24 hours of drug treatment (left panel). Responses of inflammatory genes when cells were treated first for 2 hours with the respective drug and then for 24 hours with the TNF-α (right panel). Refer to the **Materials and Methods** for abbreviations.

### Identification of three ADA drugs as RA drug candidates

Interestingly, a significant proximity to RA was found in three ADAs – GCZ, MFOM, and STGT, indicating their potential as RA drug candidates. To validate if GCZ, MFOM, and STGT can be repurposed to treat the RA, *in-vitro* experiments were performed on mouse fibroblast NIH-3T3 cells by quantifying the expression of six RA associated inflammatory genes RANTES, IL-6, CXCL-1, IL-1β, IκB and MCP-1, which are the key genes induced in the joints of RA patients [40]. We found that the computationally identified ADAs-GCZ, MFOM, and STGT were repressing the expression of these inflammatory genes. We found this effect to be similar to current commercial anti-RA drugs, methotrexate and sulfasalazine (positive control). Cimetidine, a non-inflammatory drug, appears to have no effect on the expression of inflammatory genes (negative control). TNF-α, a potent inducer of inflammatory genes, actively induced the expressions of inflammatory genes (**Fig 2C** and **Materials and Methods**).

### Proximity between DMARDs and pathways

To investigate further the network-based mechanism of drug action, we examined the pathways that are proximal to DMARDs using the gene sets of KEGG and Reactome pathways. **Fig 3A** depicts the heatmap analysis for the RA-relevant pathways. MTX was found to be located close to MAPK-, P53-, PI3K-Akt-, TGFβ-, and WNT signaling pathways. HCQ is adjoined to TLR signaling pathways, which is expected given that the target molecules are TLR7 and TLR9, but is also close to NF-κ- and VEGF signaling pathways. Type I interferon and VEGF signaling pathways in proximity to SSZ and LFNM is adjacent to a broad range of pathways. TFCN was contiguous to type I and II interferon pathways, in which JAK1 and JAK2 are the key mediators. Despite TNF inhibitors, ADLM and ETAN showed strong proximity to Fcε RI-, Jak-STAT-, and NF-κB signaling pathways rather than TNF signaling pathways.

**Fig 3.**
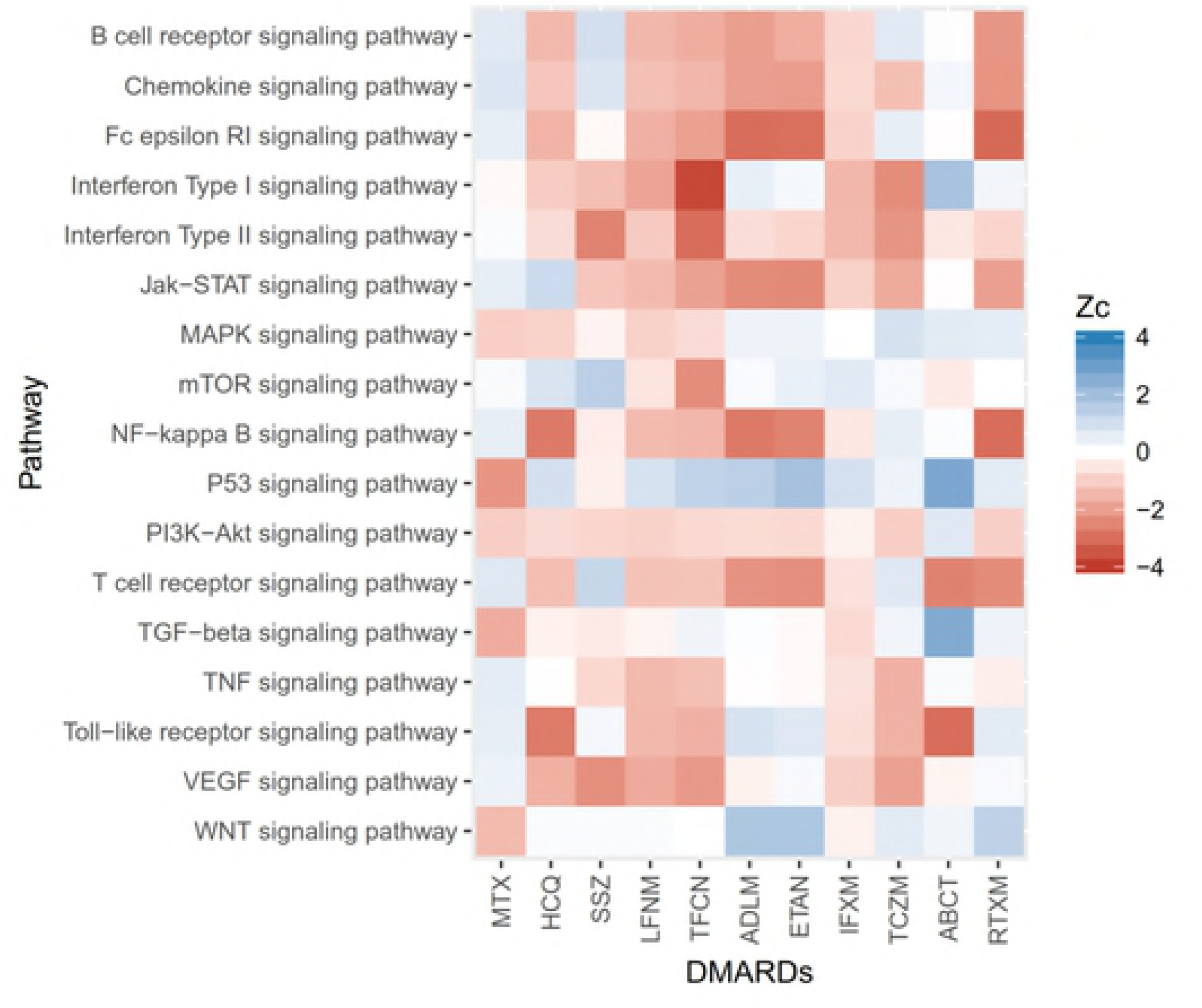
Pathway analysis of DMARDs. DMARDs and RA-relevant pathways based on proximity metrics. Refer to the **Materials and Methods** for abbreviations.

A large size of collateral targets other than TNF might dilute the direct effect of drugs on TNF signaling pathway since network-based proximity formula equivalently integrates all the distance between target proteins and a set of genes in interest. At a close range for TNF signaling pathway was IFXM, as regarded to target uniquely TNF. TCZM, ABCT, and RTXM were placed in the immediate vicinity of IL-6-related (chemokine- and interferon-), T cell receptor-, and B cell receptor signaling pathways in conformity with their specific target(s).

MTX is an anchor drug of widespread use and known clinical efficacy. However, the mechanisms by which low-dose methotrexate exerts its therapeutic effect in RA remain only partially understood and cannot be explained simply by folate antagonism [41]. To gain a better insight on the MTX-involved pathways, we measured the pathway proximity by including MTX-interacting molecules provided in other sources (DGIdb, PharmGKB, and STITCH) [42-44] as a set of targets (**S1 Fig**). MAPK-, P53-, PI3K-Akt-, TGFβ-, and WNT signaling pathways were the proximal pathways in common across the sources. In PharmGKB and STITCH data providing more adjunctive interactors, TNF, interferon type I and II signaling pathways were identified to approximate to MTX.

### Disease module identification

Known disease-associated genes tend to be investigated more extensively, which might introduce a bias and be incomplete to fully explain the pathogenesis of disease. To ease this limitation, we made the disease module based on the established RA-associated genes. A cluster of 85 highly interconnected seed genes was observed on the interactome of the 252 RA associated genes from two sources [25, 26] shown in **Fig 2A**, which we call the “proto-module”. The terms most strongly enriched within the largest connected component (LCC) of the proto-module were T cell receptor signaling pathway (*p*-value = 3.29×10^−9^), positive regulation of T cell proliferation (*p*-value = 1.89×10^−8^), positive regulation of transcription from RNA polymerase II promoter (*p*-value = 2.73×10^−8^), humoral immune response (*p*-value = 3.87×10^−7^), positive regulation of smooth muscle cell proliferation (*p*-value = 5.27×10^−7^), positive regulation of gene expression (*p*-value = 7.21×10^−7^). The rest of the seed proteins either formed small clusters (2,3 and 11 genes) or they had no interaction registered (about 54.4% of the seed proteins, 137/252).

Proteins that are involved in the same disease tend to interact with each other [13]. We prioritized the putative RA-relevant genes forming the disease module on the interactome using the DIAMOnD algorithm [33, 34] (**Fig 4A-C** and **S3 Table**). To define the boundary of the disease module, we analyzed the additional enrichment of DIAMOnD genes at each iteration using four different RA-relevant validation data - RA gene expression data, gene sets from KEGG, Reactome pathways, and GO biological process terms (**Fig 4B**). **Fig 4B-C** depict the *p*-values and construction of RA disease module by seed and DIAMOnD genes. The top 30 hits were listed in **Fig 4D**. Three of JAK kinase family genes (JAK3, JAK1, and JAK2) and the SYK gene, the emerging small molecule targets of RA, were incorporated early by priority into DIAMOnD gene although those were not assigned as seed genes. The RA disease module by the KEGG pathways was most strongly enriched with TNF signaling pathway (*p*-value = 2.08×10^−35^), followed by Fc epsilon RI signaling pathway (*p*-value = 2.92×10^−29^), Toll-like receptor signaling pathway (*p*-value = 1.02×10^−24^), and VEGF signaling pathway (*p*-value = 2.20×10^−24^).

**Fig 4.**
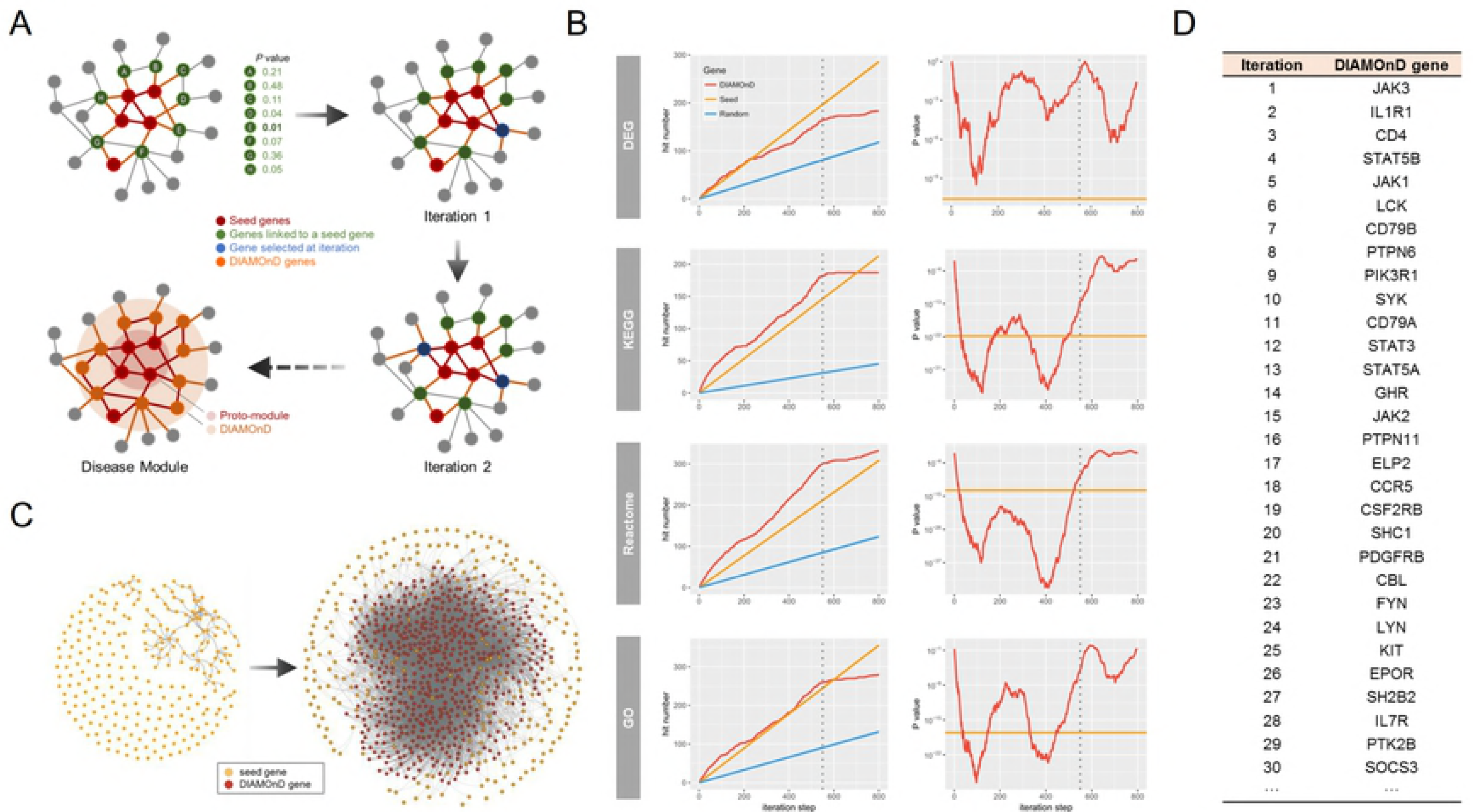
Disease module identification and validation. (**A**) Schematic network configuration from seed genes to the RA disease module. Seed genes (red) and their neighbor nodes (green) are identified iteratively. The proto-module consists of the seed genes only, while the final disease module incorporates all genes identified by the DIAMOnD method. (**B**) Validation of the identified genes through cross-reference with orthogonal data sources. The left column depicts the number of DIAMOnD genes found in the different validation datasets (RA gene expression data, gene sets from KEGG, Reactome pathways, and GO biological process terms), with the corresponding statistical signi?cance (p-value) shown on the right column. No significant gain was noted beyond 550 iteration steps (vertical gray dot line). In right column, a sliding-window approach was used to adjust for the dependence of *p*-values on the underlying set size. (**C**) The 252 seed genes and the resulting network once the 486 DIAMOnD-identified genes are added to create the 738-gene RA disease module. (**D**) List of the DIAMOnD genes at the first 30 iterations.

### Perturbation of disease modules by DMARDs

To comprehend the pathways or cellular processes guided by the disease module, we performed the functional enrichment analysis for all seed and DIAMOnD genes using the DAVID software [35] and constructed the enrichment map. Graph of enriched GO terms of biological process was made of 120 nodes with 488 edges and the LCC was comprised of 107 nodes (**Fig 5A** and **S2 Fig**). To check how close a drug was to each node enough to perturb the process, we calculated the network-based proximity of drug targets to gene set of each node in the same way for disease and pathway proteins. We regarded the node significantly proximal to a certain drug (*z*_*c*_ < −0.15) as being perturbed.

**Fig 5.**
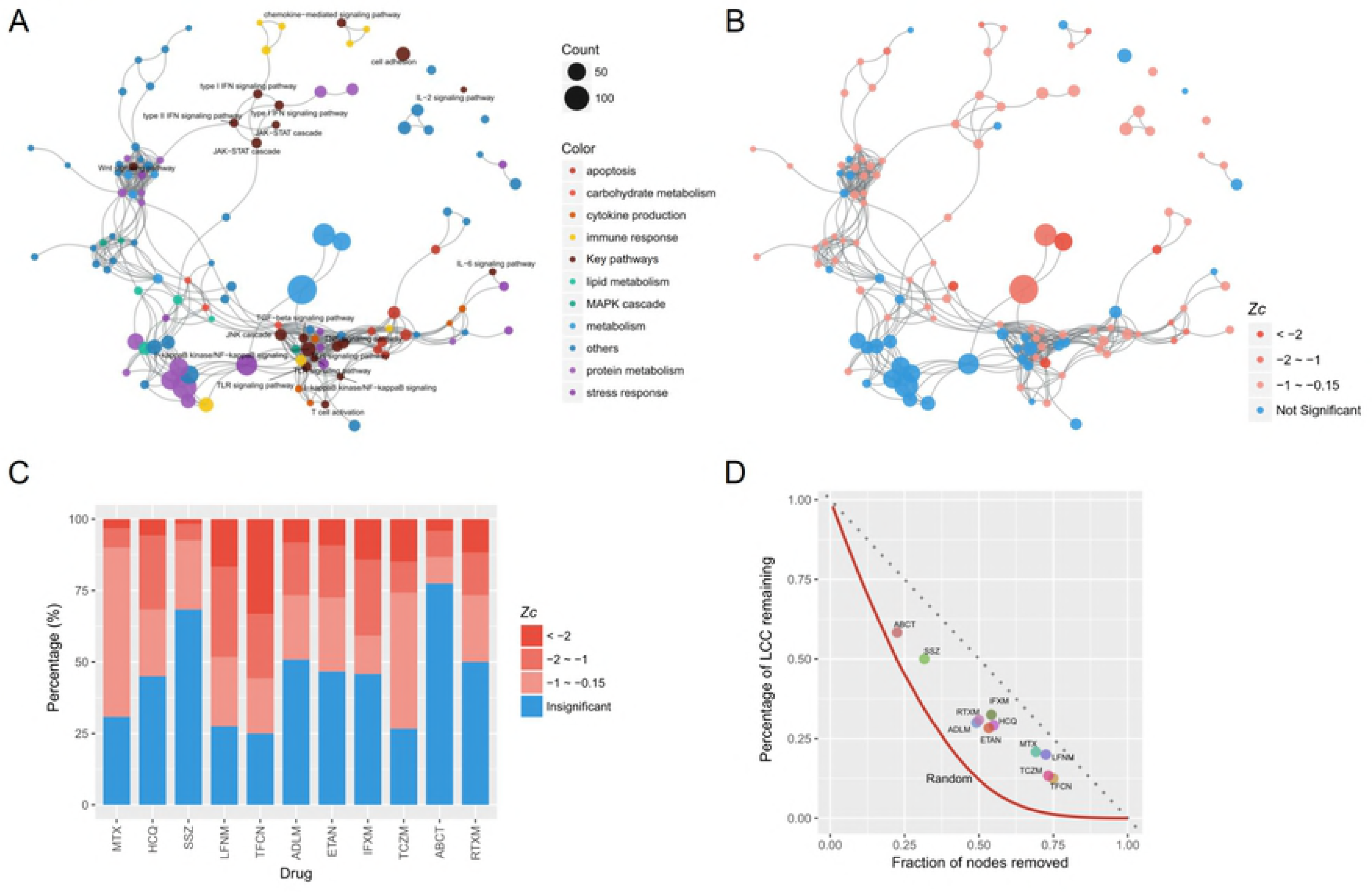
RA disease module perturbation by DMARDs. (**A**) Enriched pathway map of the RA disease module. Nodes represent distinct GO terms. Node size is proportional to the number of genes in the respective GO term. Edge was determined by the Jaccard similarity coefficient (edge cut-off is 0.1 distance). (**B**) Perturbation of the disease module by the MTX drug. Nodes significantly proximal to drug targets are red, insignificant nodes are blue (*z* ≤ –0.15). (**C**) Ratio of nodes based on the proximity between the DMARDs and the disease module provide the specificity and generalization of the respective DMARD. (**D**) The percentage of the largest connected component (LCC) that remains when treated by DMARDs (points) or under random node failure (red line, 10^3^ simulations). Refer to the **Materials and Methods** for abbreviations.

In an attempt to understand the action mechanisms of the respective drugs, we analyzed the biological processes and molecular function of the GO terms in its disease network. The terms most proximal to MTX were ‘positive regulation of protein insertion into mitochondrial membrane involved in apoptotic signaling pathway’ (*z*_*c*_ = –2.41), ‘DNA damage response, signal transduction by p53 class mediator resulting in cell cycle arrest’ (*z*_*c*_ = –2.26), ‘rhythmic process’ (*z*_*c*_ = –2.11), and ‘transcription from RNA polymerase II promoter’ (*z*_*c*_ = –2.04). MTX perturbed 83 (69.2%) nodes, which were colored with red hue in **Fig 5B**. Less nodes (66, 55%) and smaller overlap with MTX was found in the case of HCQ. LFNM and TFCN were found to have a high overlap with MTX. It is noteworthy that TFCN attacked the largest size of nodes (90, 75.0%) and was the closest in proximity to 40 nodes (33.3%) with *z*_*c*_ < –2. The distribution of perturbed nodes by each DMARD is provided in **Fig 5C** and **S3 Fig**.

In a network context, robustness refers to the system’s ability to carry out its basic functions even at the absence of some network nodes or links. The connectivity of LCC is regarded as the backbone of robustness [37]. To investigate the robustness of the disease module network, we computed the evolution of the size of the LCC under random failures of network nodes [36, 37] (**Fig 5D**). Our results (1000 simulations) argue that half of the network could collapse when at least 22% of the nodes are disturbed on average. All DMARDs, except for ABCT, impacted over a quarter of the nodes and collapsed over half of the disease module network. TFCN and TCZM were found to confer the highest robustness with respect to network stability, while ABCT had the weakest stability of all.

### Module-driven validation of network-based proximity by tocilizumab

To test the applicability of network-based proximity by DMARDs, we imported the gene expression dataset of RA synovial tissues, which was obtained from 12 RA patients on two separate occasions before and after a 12-week TZCM treatment (GSE45867) [45]. To compare the paired samples before and after TZCM treatment, ssGSEA was used to generate enrichment scores for gene sets of GO terms by RA disease module. Significant change (*p*-value < 0.05) was found in 49 GO terms and the 10 main signaling pathways or cytokine-related process of reduced scores are identified (**Fig 6A** and **S4 Fig**). which are in agreement with the pathway proximity result of TCZM (**Fig 3A**). We applied the outcome to a binary classification (labels of “good” and “not-good” responder) by EULAR response criteria and 6 patients were classified as “good” responders. As compared between “good” and “not-good” responder, score decrease in the regulation of phosphatidylinositol 3-kinase signaling (GO:0014066) and ERK1 and ERK2 cascade (GO:0070371) were significant in the “good” responder category as opposed on the “not-good” one (*p*-value < 0.05; **Fig 6B**).

**Fig 6.**
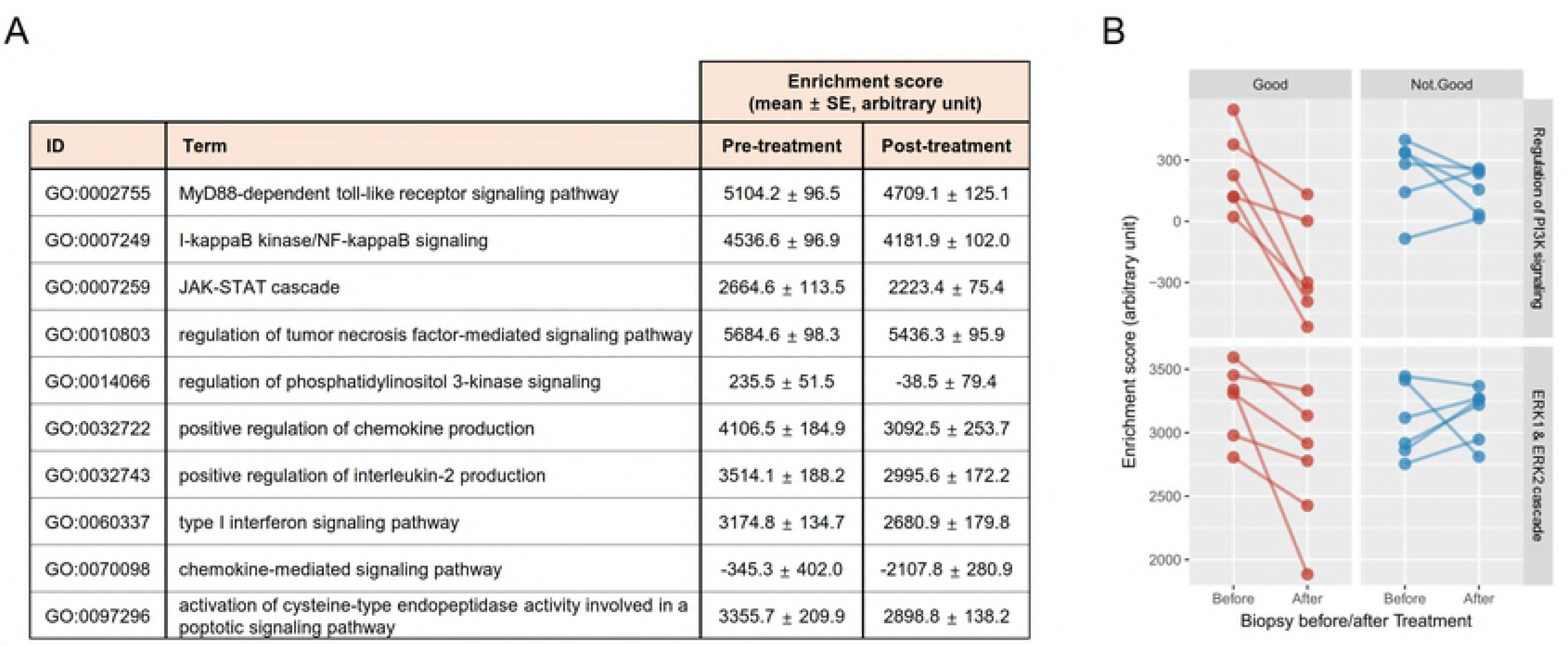
Validation of network-based proximity by tocilizumab. ssGSEA was performed for paired synovial biopsy samples obtained from naïve RA patients before and 12 weeks after TCZM treatment (GSE45867) [45]. Comparisons between the groups were made using paired or unpaired *t*-test. (**A**) Ten main signaling pathways or cytokine-related process of significant reduction in enrichment scores after treatment (*p*-value < 0.05). (**B**) Change of enrichment score in ‘regulation of phosphatidylinositol 3-kinase signaling’ and ‘ERK1 and ERK2 cascade’ by outcome. Score decrease in “good” responders is significant as compared with “not good” responders (*p*-value < 0.05).

### Proximity between DMARDs and side effect proteins

Network-based proximity does not always imply that the drug will assuage the corresponding disease. The drug could negatively affect the nearby molecules of normal physiology, leading to unwanted side effects [31]. To see whether the known side effects of DMARDs could be obtained by extrapolating from the network-based analysis, we checked the proximity of chemical DMARDs and the three candidate repurposing ADA drugs (GCZ, STGT and MFOM) to the protein sets predicted to induce the side effects, as shown in **Fig 7**. Since biologic DMARDs of synthetic immunoglobulin-like forms are deemed to interact with the defined targets and to be unable to pass through cell membrane, they were excluded for this analysis. In the case of MTX, 20 of the 64 proteins significantly proximal to MTX (31.3%) have at least one corresponding side effect that is listed in the side effects of MTX from the SIDER database (**S4 Table**). For HCQ, 7 proteins provoking at least one SIDER side effect were found in the proximal range with some of them associated with visual disturbance (CACNA1F, RHO), myopathy (RYR1), and dizziness (CACNA1A), hence need more attention during clinical practice (**S2 Table**). Only 2 proteins having SIDER side effects were observed in TFCN. This might be because its side effects were not sufficiently surveyed due to their short experience in clinical practice. There was no protein having the corresponding side effects in the outside of proximal area except for one (ATP1A2) in LFNM.

**Fig 7.**
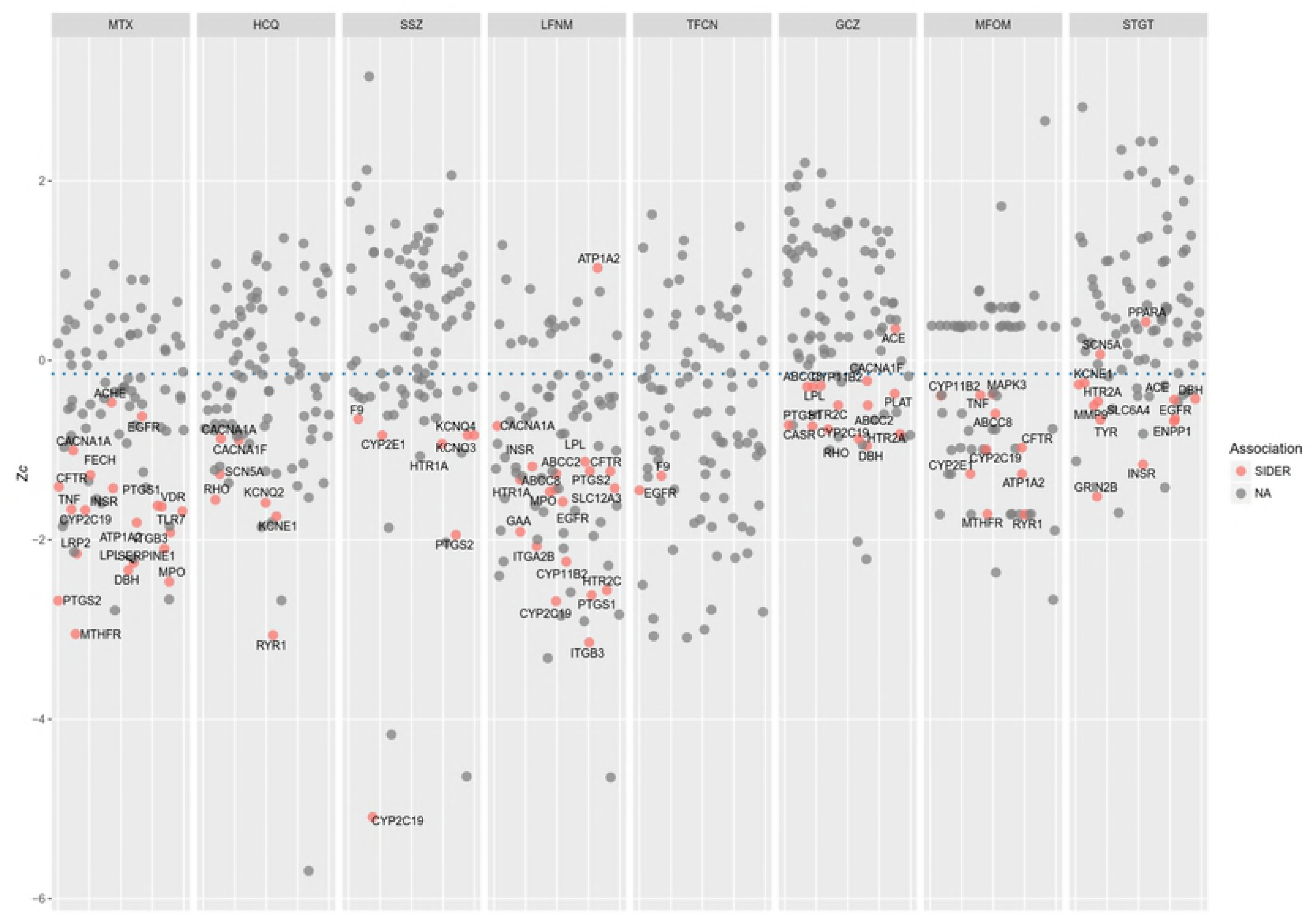
Proximity between DMARDs and side effect proteins. Proteins eliciting the side effects listed in SIDER database by drugs were marked as red dots [32]. A proximity of *z* ≤ –0.15 (blue dot line) is regarded to be significantly proximal to drug [8]. Refer to the **Materials and Methods** for abbreviations.

## Discussion

As expected, most of DMARDs were found to be significantly proximal to RA-associated genes. Determination of the action mechanism of low-dose MTX for the observed clinical effects in RA has been quite challenging, despite MTX being an anchor drug, [41]. Interestingly, we found that antagonism of folate-dependent processes, the defined mechanism of MTX, was closely linked to MAPK-, P53-, PI3K-AKT-, TGF-β-, and WNT signaling pathways, something that is corroborated by the interaction of MTX with the MAPK-, P53-, or WNT signaling pathways depending on cell types and experimental condition [46-48]. A significant finding of our analysis is the identification of three ADAs (GCZ, MFOM, and STGT) as candidates for drug repurposing in the case of RA treatment. Subsequent *in-vitro* experiments on mouse fibroblast NIH-3T3 cells proved the engagement of GCZ, MFOM, and STGT in the repression of six RA related inflammatory genes. We compared the proximity of DMARDs with those of controls including ADA and statins. It is intriguing that GCZ, MFOM, and STGT stand in a significant proximity to a group of RA-associated genes. In the literature, GCZ has an anti-inflammatory effect by inhibiting monocyte adhesiveness to vascular cells through reduction of oxidized low-density lipoprotein and advanced glycation end product [49]. MFOM showed a suppressive effect on TNF-α–dependent IκB degradation and expression of proinflammatory mediators IL-6 and IL-1β [50]. Moreover, the anti-arthritic effect of MFOM was recently demonstrated in collagen-induced arthritis model [51]. DPP4 has capability to directly activate the MAPK and NF-kB signaling cascade [52] and DPP4 inhibitors showed additional anti-inflammatory and immunomodulatory effects in autoimmune diabetes or myocarditis animal models by recovering the number of CD4+CD25+Foxp3+ cells or inhibiting expression of IL-2, TNF-α and IL-6 [53, 54]. Since these drugs might be beneficial to ease off the inflammatory response, they could be considered by priority in case of a patient having both RA and diabetes unless contraindicated. These results suggest that network-based proximity inform us of candidate drugs to be repurposed in other disease [8].

The networks of the RA-associated genes are sparse and fragmented probably due to inherent incompleteness of interactome, false positives, and missing disease genes bridging between seed genes [25, 26]. We further reconstituted the RA disease module of 738 genes from 252 seed genes (proto-module), with a distinct signature: while the proto-module largely focused on the T-cell receptor signaling pathway, the RA disease module was enriched with a list of RA-relevant signaling pathways and its functional enrichment analysis was in agreement with the known RA pathophysiology. In parallel, HCQ and LFNM have a better proximity to RA disease module rather than just seed genes and the emerging small molecule targets such as JAK family and SYK were captured at early stages.

We investigated the size and range of disease module differentially covered by each DMARD to understand its mode of action and topology in the network-based framework. We noticed a variable size and differential coverage of the disease network by the drugs. When a patient is first diagnosed with RA, MTX monotherapy is usually used as the first line of defense [55, 56]. If the patient is refractory to MTX, HCQ and/or SSZ are added to the regimen. Our analysis argues that the combination of HCQ and/or SSZ with MTX could create a good synergy effect because of their complementarity in the LLC network (**S3** and **S5 Fig**).

Our analysis shows that if MTX-based double or triple therapy fails, the regimen can be switched to LFNM or TFCN, as these are suitable substitutes given the more extensive and less overlapping to MTX sub-network they perturb, including all the critical hubs. It is well-known that efficacy of TNF inhibitors is enhanced by their combination with MTX [57]. This is inferable by the unshared subnetwork and complementary relationship between TNF inhibitors and MTX. It is also plausible that TCZM monotherapy produces a similar effect to combination of TNF inhibitor and MTX [57] thanks to the merged coverage by TNF inhibitor and MTX. ABCT inhibits T cell activation, selectively blocking the specific interaction of CD80/CD86 receptors to CD28 and covered the least territory of RA disease module, resulting the weakest direct effect to collapse the disease module. Indeed, ABCT is criticized for being slower in reducing disease activity although the final outcome is equivalent to other biologic DMARDs [58]. However, ABCT was estimated to be a best proximity to RA-associated genes (seed genes) and this is probably because T-cell receptor signaling pathway is the main enrichment of RA-associated genes. Taken together, the disease module and the proximity metric can be used together to inform how to develop better treatment strategies that rely on drug combinations. In effect, when applied to real gene expression data, pathways proximal to TCZM were in accordance with down-regulation (**Fig 3** and **6**), with the PI3K-AKT pathway and ERK cascade might be contributing factors determining the responsiveness for their different transition by outcome.

We found that the proximity between drug targets and proteins that elicit side effects is a suitable metric for the probability of side effects in the case of DMARDs. It is noteworthy that information on retinal toxicity and myopathy was captured in the case HCQ, as these are rare but serious complications that often lead to patient death. SIDER proteins in DMARDs were more proximal as compared with those in 3 candidate ADAs (mean *z*_*c*_ –1.64 versus –0.58, *p*-value < 0.001), possibly because of the difference in the main target molecules (inflammatory/immunologic versus metabolic). Although, the proximity metric provides a useful quantification of risk, it is just one of the parameters that should be taken into account, as many proteins that are not directly associated with side effects are impacted by these drugs, hence indirect effects are likely.

Our results argue that network-based drug-disease proximity offers a novel perspective to a drug’s therapeutic effect. In this work, we investigated just one of the dimensions that need to be taken into account when determining potential drug candidates for each case, as the actual clinical outcome is determined by a plethora of variables, including dosage, drug interactions, comorbidity, metabolic capacity to drugs, genetic susceptibility, among others. Still, the network-based approach that we describe here provides a fast and efficient way to determine likely candidates for drug repurposing and understand their underlying mechanisms, with far-reaching applications to various diseases beyond RA.

## Acknowledgements

We would like to thank Prof. Henry Ho for providing us the mouse fibroblast NIH-3T3 cell line, and Prof. Alyssa Panitch for sharing the animal cell culture facility.

## Author Contributions

**Conceptualization:** Ki-Jo Kim, Ilias Tagkopoulos

**Data curation, formal analysis, & investigation:** Ki-Jo Kim, Navneet Rai, Minseung Kim

**Methodology:** Ki-Jo Kim, Navneet Rai

**Visualization:** Ki-Jo Kim, Navneet Rai, Minseung Kim

**Supervision:** Ilias Tagkopoulos

**Writing – original draft:** Ki-Jo Kim

**Writing – review & editing:** Navneet Rai, Ilias Tagkopoulos

## Supporting Information

**S1 Fig. Heatmap of proximity between MTX-interacting molecules from different sources (DrugBank, DGIdb, PharmGKB, and STITCH) and RA-relevant pathways.** MTX-interacting molecules from each source were listed in the upper panel.

**S2 Fig. Functional enrichment map using DAVID for combination of RA-associated genes and DIAMOnD genes.** Same as **Figure 5A** except for detailed annotation.

**S3 Fig. Network perturbation by proximity between all DMARDs and disease module.** Layout of the network is the same as the enrichment map of RA disease module (**Figure 5A**). Nodes significantly proximal to drug targets are colored by red hue depending on the degree of proximity and insignificant nodes are coated by blue color.

**S4 Fig. Ten main signaling pathways or cytokine-related process of significant reduction in enrichment scores after treatment (*p*-value < 0.05).**

**S5 Fig. Percentage of the shared, distinct, or united nodes on RA disease module by DMARDs.** Each column is the baseline DMARD and rows are the counterparts in comparison. Values were calculated from the number of the corresponding nodes divided by the total number of nodes on RA disease module (n=120).

**S1 Table. 252 RA-associated genes from PheGenI and DisGeNET.**

**S2 Table. Information on drug-target molecules from the DrugBank database.**

**S3 Table. 550 genes from DIAMOnD algorithm.**

**S4 Table. Proximity between DMARDs and side effect proteins.**

